# Single-cell and nucleus RNA-seq in a mouse model of AD reveal activation of distinct glial subpopulations in the presence of plaques and tangles

**DOI:** 10.1101/2021.09.29.462436

**Authors:** Gabriela Balderrama-Gutierrez, Heidi Liang, Narges Rezaie, Klebea Carvalho, Stefania Forner, Dina Matheos, Elisabeth Rebboah, Kim N. Green, Andrea J. Tenner, Frank LaFerla, Ali Mortazavi

**Affiliations:** University of California, Irvine, Department of Developmental and Cell Biology, Irvine, CA 92697, USA; University of California, Irvine, Institute for Memory Impairments and Neurological Disorders, University of California, Irvine, CA 92697, USA; University of California, Irvine, Center for Complex Biological Systems, Irvine, CA 92697, USA; University of California, Irvine, Department of Neurobiology and Behavior, University of California, Irvine, Irvine, CA 92697, USA; University of California, Irvine, Department of Molecular Biology and Biochemistry, Irvine, CA 92697, USA

## Abstract

Multiple mouse models have been generated that strive to recapitulate human Alzheimer’s disease (AD) pathological features to investigate disease mechanisms and potential treatments. The 3xTg-AD mouse presents the two major hallmarks of AD, which are plaques and tangles that increase during aging. While behavioral changes and the accumulation of plaques and tangles have been well described in the 3xTg-AD mice, the subpopulations of neurons and glial cells present throughout disease progression have not been characterized. Here, we used single-cell RNA-seq to investigate changes in subpopulations of microglia, and single-nucleus RNA-seq to explore subpopulations of neurons, astrocytes, and oligodendrocytes in the hippocampus and cortex of aging 3xTg-AD as well as 5xFAD mice for comparison. We recovered a common path of age-associated astrocyte activation between the 3xTg-AD and the 5xFAD models and found that 3xTg-AD-derived astrocytes seem to be less activated. We identified multiple subtypes of microglia, including a subpopulation with a distinct transcription factor expression profile that showed an early increase in *Csf1* expression before the switch to disease associated microglia (DAM). We used bulk RNA-seq in the hippocampus of 3xTg-AD mice across their lifespan to identify distinct modules of genes whose expression increases with aging and worsening pathology. Finally, scATAC-seq revealed multiple subpopulations of cells with accessible chromatin in regions around genes associated with glial activation. Overall, differences between the main glial groups point to a slower activation process in the 3xTg-AD model when compared to the 5xFAD. Our study contributes to the identification of progressive transcriptional changes of glial cells in a mouse model that has plaques and tangles, thus providing information to aid in targeted AD therapeutics that could translate into positive clinical outcomes.

## INTRODUCTION

Alzheimer’s disease (AD) is a progressive neurodegenerative disorder characterized by inflammation and accumulation of extracellular amyloid beta (Aβ) plaques and neurofibrillary tangles, culminating with neuronal loss and brain tissue shrinkage ^1^. The resulting brain tissue damage over time causes cognitive impairment and memory loss ^2^. Uncovering the underlying mechanisms that lead to tissue loss and memory impairment remains a challenge. Several clinical trials have investigated drug candidates to stop or reverse AD progression, but the failure rate of these trials has been extremely high ^3^. Animal models play an essential role in preclinical studies to help understand disease mechanisms and test possible therapeutic interventions. However, promising findings from mouse models do not always translate into successful clinical trials.

Many mouse models of AD have been developed using transgenes that overexpress human APP alongside early onset of AD (EOAD) and familial AD mutations, rather than modelling sporadic/late onset AD (LOAD) that constitute the vast majority of AD cases ^4^. Most of these models only develop one of the two main AD hallmarks, which are amyloid plaques. Hence, they fail to recapitulate the pathology observed in humans in several ways. One extensively characterized model is the 5xFAD mouse that develops an aggressive plaque pathology marked by early amyloid peptide accumulation at around 2 months of age (the equivalent of an 18-year-old human) but no tangles ^5,6^. Thus, 5xFAD is a valuable model for amyloidosis but does not fully recapitulate AD. Another commonly used model is the 3xTg-AD mouse (Tg(APPSwe,tauP301L)1Lfa Psen1tm1Mpm/Mmjax), which presents the combination of both plaques and tangles and is considered a more complete mouse model of AD ^7,8^. The 3xTg-AD mouse model develops its pathological features more slowly throughout its lifespan compared to other AD models and thus may be useful to study the impact aging has on the phenotype. Deep phenotyping and characterization of the 3xTg-AD mouse has been explored in a companion publication ^8^.

Glial cells play an important role throughout central nervous system (CNS) development, disease, and homeostasis ^9^. Besides neurons, glial cells are key players in several neurodegenerative diseases, including AD ^10,11^. Glial cells show functional diversity and distinct transcriptional responses to the presence of plaques and tangles in the brain ^12,13^. GWAS studies in populations of patients with LOAD identified risk loci associated with microglia-specific genes, such as *TREM2*, and, *HLA-DQA1* ^14,15^. Microglia are the resident macrophages of the brain. They promote immunosurveillance of the brain and help maintain homeostasis. However, in the inflammatory AD brain, microglia can assume a disease enhancing phenotype (a subset of disease associated microglia termed DAM) showing morphological changes, as well as upregulation of genes such as *Cst7, Trem2* and *Tyrobp* ^16^. Multiple classifications have been proposed to highlight microglial states in brain homeostasis and injury, including: DAM, homeostatic microglia, proliferative-region microglia, and variations of DAMs ^16–18^. Each of these microglial subtypes displays a unique transcriptional signature.

Astrocytes also present distinct activation states, including states associated directly with AD, whereby they show upregulation of several genes ^19,20^. *CLU*, also known as *APOJ*, is a glial gene associated with LOAD and belongs to the same apolipoprotein family as *APOE. CLU* is expressed in neurons and astrocytes of human and mouse, but its expression is much higher in astrocytes, especially after injury ^21^. *Clu* is one of the first markers of astrocyte activation after injury ^22^ and its overexpression results in reduction of Aβ plaques in the *APP/PS1* mouse model of AD. *Gfap* and *Serpina3n* are also upregulated in reactive astrocytes during neurodegeneration ^20^. During AD, astrocyte activation affects its interactions with other cell types, including oligodendrocytes ^23^. The main function of oligodendrocytes is the wrapping of neuronal axons with myelin to increase the efficiency of action potentials, but they are also capable of remyelination ^24,25^. In the 5xFAD model, a decrease in astrocyte-oligodendrocyte interactions is explained by the decrease of oligodendrocyte progenitor cells (OPCs) starting at 3 months ^23^. In a similar vein, the 3xTg-AD mouse displays a loss of OPCs at 6 months of age ^23,26^, which contributes to AD pathology. 3xTg-AD mice present changes in oligodendrocyte populations even before the appearance of plaques ^27^.

Single-cell platforms that rely on microfluidic devices do not perform well with brain cells given the obstructions caused by the large size of neurons and astrocytes, as well as the myelin generated by oligodendrocytes. Consequently, single-nucleus based approaches have become an alternative to address the diversity in populations of large cells or in tissues that are resistant to dissociation ^28^. However, there are reports that single-nucleus RNA-seq misrepresents the diversity of subtypes of human activated microglia that can be detected using whole single-cells ^29^. Furthermore, most single-cell investigation of microglial subtypes have been done using whole AD mouse brains, instead of specific brain regions. In addition, studies that evaluated region-specific datasets are focused on development or early life stages ^30^. Time courses describing microglial diversity, while useful, are derived from mouse models that present only plaques, not tangles and thus do not recapitulate the human pathology ^31^. Longitudinal studies describing the progression of cell and region-specific transcriptional changes in a mouse model with plaques and tangles are still needed.

Transcriptional regulation can be inferred by analyzing the chromatin accessibility state using ATAC-seq paired with RNA-seq data. Chromatin accessibility is one of the determinants of transcription factor binding and can predict gene expression patterns ^32^. Studies in neurons derived from induced pluripotent stem cells from EOAD patients observed that chromatin patterns affect differentiation events and mirror gene expression signatures ^33^. Human and mouse microglia have shown conserved transcriptomic and epigenomic phenotypes ^34^. A recent study showed that isogenic human ESC–derived microglia harboring AD variants in the *TREM2* locus present an “AD-primed” state, including changes at chromatin levels ^35^. The investigation of chromatin accessibility patterns during microglial activation and cellular predisposition to brain injury can deliver further insights to AD studies.

Here, we present the first characterization of the transcriptome of subpopulations of cells derived from the cortex and from hippocampus throughout the lifespan of the 3xTg-AD mouse model of AD as part of our single-cell and single-nucleus RNA-seq pipeline developed for MODEL-AD (https://www.model-ad.org) ^36^. We applied a time course of bulk RNA-seq in the hippocampus of 3xTg-AD mice at 4-, 12- and 18-months of age. We identified modules related to the immune system, synaptic activity, and myelination, which we compared to existing gene sets in human AD and the 5xFAD mouse model. We applied a single-nucleus RNA-seq (snRNA-seq) time course in cortex and hippocampus of 12-, 18-, and 24-month-old 3xTg-AD mice and identified distinct populations of neurons, astrocytes, and oligodendrocytes that are present during the late stages of neurodegeneration. We also performed single-cell RNA-seq of microglia at 12-, 18-, and 24 months of age in order to identify activated subtypes, given the low capture of microglia in our snRNA-seq dataset. Finally, we assayed the chromatin profiles of microglia from the cortex of a 24-month female 3xTg-AD mouse using single-cell ATAC-seq, in order to identify the chromatin accessibility profile for activated microglia.

## RESULTS

### Gene expression changes in the 3xTg-AD mouse model of AD suggest increased activation of microglia and astrocyte during progression of pathology

We analyzed a bulk RNA-seq time course of 3xTg-AD hippocampus at 4, 12, and 18 months of age to understand the transcriptional changes that occur during aging in this model. Female 3xTg-AD (as well as female 5xFAD) mice have shown increased pathology when compared to males, likely due to the *Thy1* mini-gene, which regulates the expression of the cDNAs in the transgenes ^6^. Therefore, we expected stronger upregulation of AD-associated genes in females. We only detected higher expression of known DAM markers such as *Cst7* in females. Since the goal of this study is to elucidate pathways involved in AD, we focused the rest of our bulk analysis on females. We identified 3 outliers (∼0.6 Pearson) that were excluded from further analysis (Figure S1). We applied Weighted Gene Correlation Network Analysis (WGCNA) to determine the correlation patterns among genes expressed in the hippocampus of the aging 3xTg-AD mice (Figure 1A). We identified 15 distinct modules, which we correlated with age, genotype, and modules previously described in the 5xFAD model ^6^. The paleturquoise, darkolivegreen, and magenta modules presented the highest correlation with the 3xTg-AD genotype (Figure 1B). Of these, the paleturquoise showed the strongest correlation (p-value 0.03) and consists of 111 genes whose expression increased with age. The paleturquoise module contained genes associated with inflammation and activated microglia and astrocytes, such as *Cst7, Ctss, Tyrobp, Gfap*, and *Trem2* (Figure 1C) ^20,37,38^. The darkolivegreen module contains a set of 93 genes (Figure 1D) known to be highly expressed in neurons including *Slc17a6, Rora*, and *Adra1b* ^39^. The expression of these genes decreased with age in the 3xTg-AD mice, suggesting a possible gradual neuronal loss in this model, which agrees with the previously described loss of parvalbumin-positive interneurons in the subiculum of 18-month-old 3xTg-AD females ^8^. The magenta module has 409 genes whose expression peak at 18 months (Figure 1E). This module contains myelination-associated genes *Mal, Myrf, Sox10, Cspg4/Ng2*, and *Plp1*^40^. Our results suggest an age-dependent increase in inflammation and activation of microglia and astrocytes.

**Figure 1.**
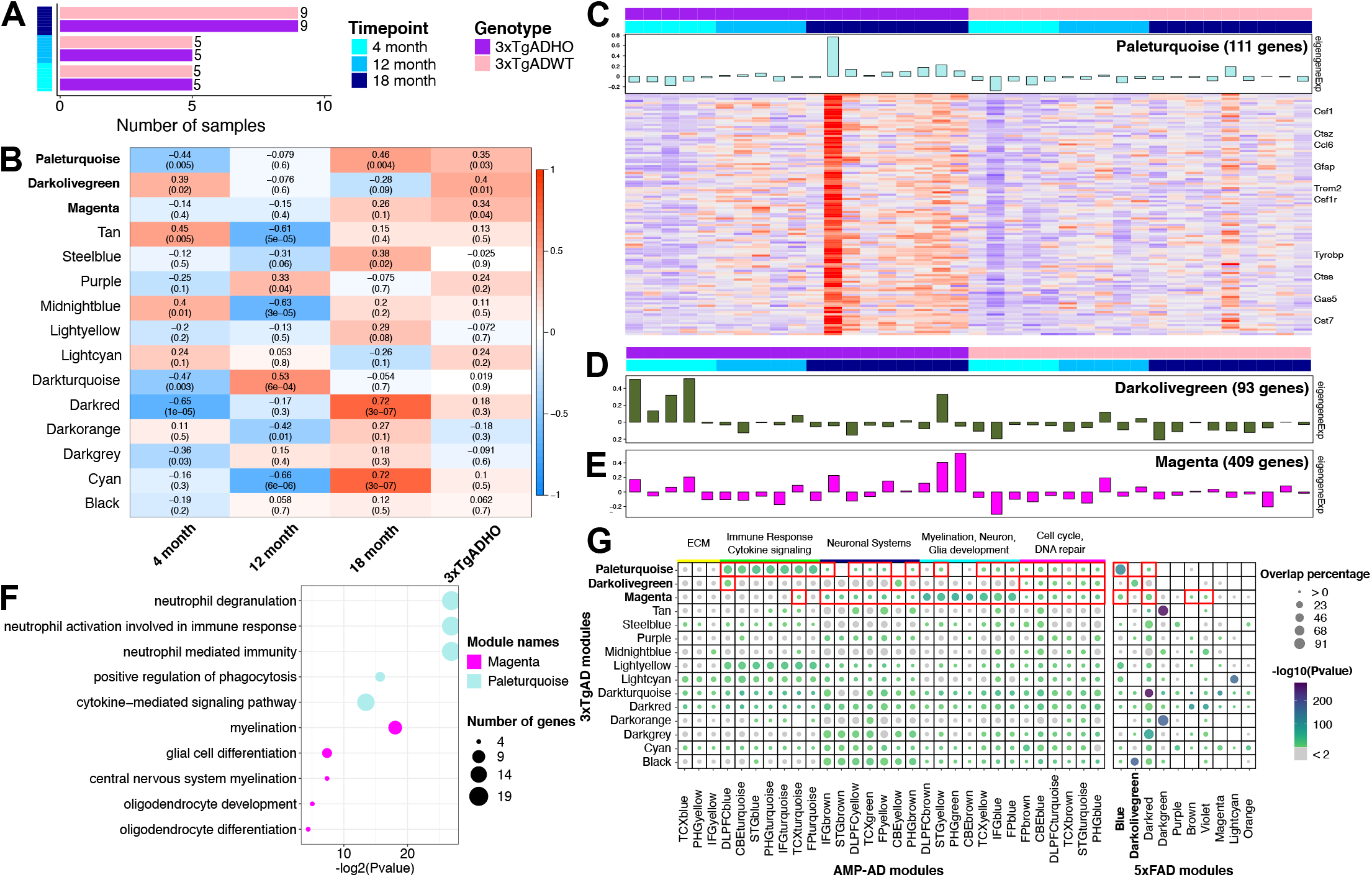
Bulk RNA-seq analysis reveals multiple 3xTg-AD gene modules that are also enriched in human AMP-AD modules. A) Number of samples assayed for each type of mouse model (3xTg-AD and Wild type). B) Module Trait Relationships (MTRs) correlation heatmap and corresponding p-values between the detected modules (y-axis) and sample features (x-axis). C-E) Bar plot of eigengene expression across samples and heatmap of gene expression matrix in C) paleturquoise module, D) darkolivegreen module, and E) magenta module. F) Significant gene ontology (GO) (adjusted p-value < 0.05) for magenta and darkolivegreen modules. G) Comparison of 3xTg-AD modules against AMP-AD and 5xFAD modules.

We repeated the WGCNA analysis including cortex and male samples. We did not find a significant correlation between male samples and age, tissue, or genotype, corroborating with the reduced pathology reported in male mice ^41^. In addition, cortex samples did not show correlation with the traits analyzed (Figure S2), which agrees with the lack of Thio-S plaques in cortical regions of 3xTg-AD mice ^8^. We also performed Gene Ontology (GO) analysis to identify biological processes and pathways enriched in the three 3xTg-AD-associated modules (Figure 1F). The paleturquoise module was enriched for inflammation-related GO terms. The darkoliveGreen module was enriched for one biological process, which did not pass the FDR threshold (Figure S3). The magenta module was enriched for GO terms associated with myelination processes, such as glial cell differentiation and oligodendrocyte development. While the cyan and darkred modules showed a high correlation with the latest timepoint/age, they did not show association with the 3xTg-AD genotype, suggesting that they represent aging-induced genes. In summary, we detect two distinct modules that contain genes associated with glial inflammation in 3xTg-AD females, one of which was associated with oligodendrocytic genes.

Consortiums such as the Accelerating Medicines Partnership Program for AD (AMP-AD) and MODEL-AD have described modules of genes associated with AD in humans and 5xFAD mice, respectively ^6,42^. To investigate the relevance of the gene expression changes seen in our 3xTg-AD time course to human AD and other mouse models, we compared our modules to previously identified AMP-AD and 5xFAD modules, respectively (Figure 1G). Our paleturquoise (inflammation-associated) and magenta (myelination-associated) modules showed high overlap with the AMP-AD inflammation (green) and myelination/neuron/glial development (turquoise) modules, respectively. Lower overlaps of those modules extend to neuronal systems (only magenta) and cell cycle/repair. The 5xFAD blue module is linked to inflammatory response and showed the highest overlap with our 3xTg-AD paleturquoise module, suggesting that both models share inflammatory signaling pathways ^6^.

Comparison of the 3xTg-AD darkolivegreen module with AMP-AD human modules show significant overlap with neuronal systems (CBEyellow module), as well as to cell cycle and DNA repair modules (Figure 1G). The 3xTg-AD darkolivegreen module showed a significant overlap with the 5xFAD darkolivegreen module, both of which are associated with the AD genotype and expression of those genes decrease with age. Taken together, these findings confirm that the 3xTg-AD develops a gradual age-related inflammatory response with gene expression changes corresponding to glial cell activation. In addition, the age-related decrease in expression of genes from a neuron-associated module (darkolivegreen) suggests a subtle neuronal loss that coincides with the previously described loss of PV^+^ interneurons in 3xTg-AD ^8^.

### snRNA-seq identifies distinct brain cell populations between the 3xTg-AD and 5xFAD mice

Although bulk correlation network analyses are useful to help understand the progression of the pathology in mouse models of AD, they do not fully address the cellular heterogeneity present in the brain tissues during neurodegeneration. Thus, we also performed snRNA-seq in the cortex and hippocampus of the aging 3xTg-AD mice and controls (WT) to investigate the subpopulations of cells present during late stages of neurodegeneration at 12, 18, and 24 months (Table S1). Considering that the current 3xTg-AD mice develop pathology later in life^8^, we also included 5xFAD 18-month female samples to compare the late transcriptional changes in both models. We obtained a total of 16,482 nuclei that were classified into 31 distinct clusters (Figure 2A). We used the expression of known markers of microglia, astrocytes, and neurons to identify the main populations in our data (Figure 2B & Table S2). Cluster N26 represents microglia, which are a small fraction of the cells recovered, similar to what has been observed in other single-nucleus RNA-seq studies ^29^. Astrocytes represented ∼20%, oligodendrocytes, and OPC’s ∼12%, and microglia 1% of the nuclei in our study, with the remainder being neurons.

**Figure 2.**
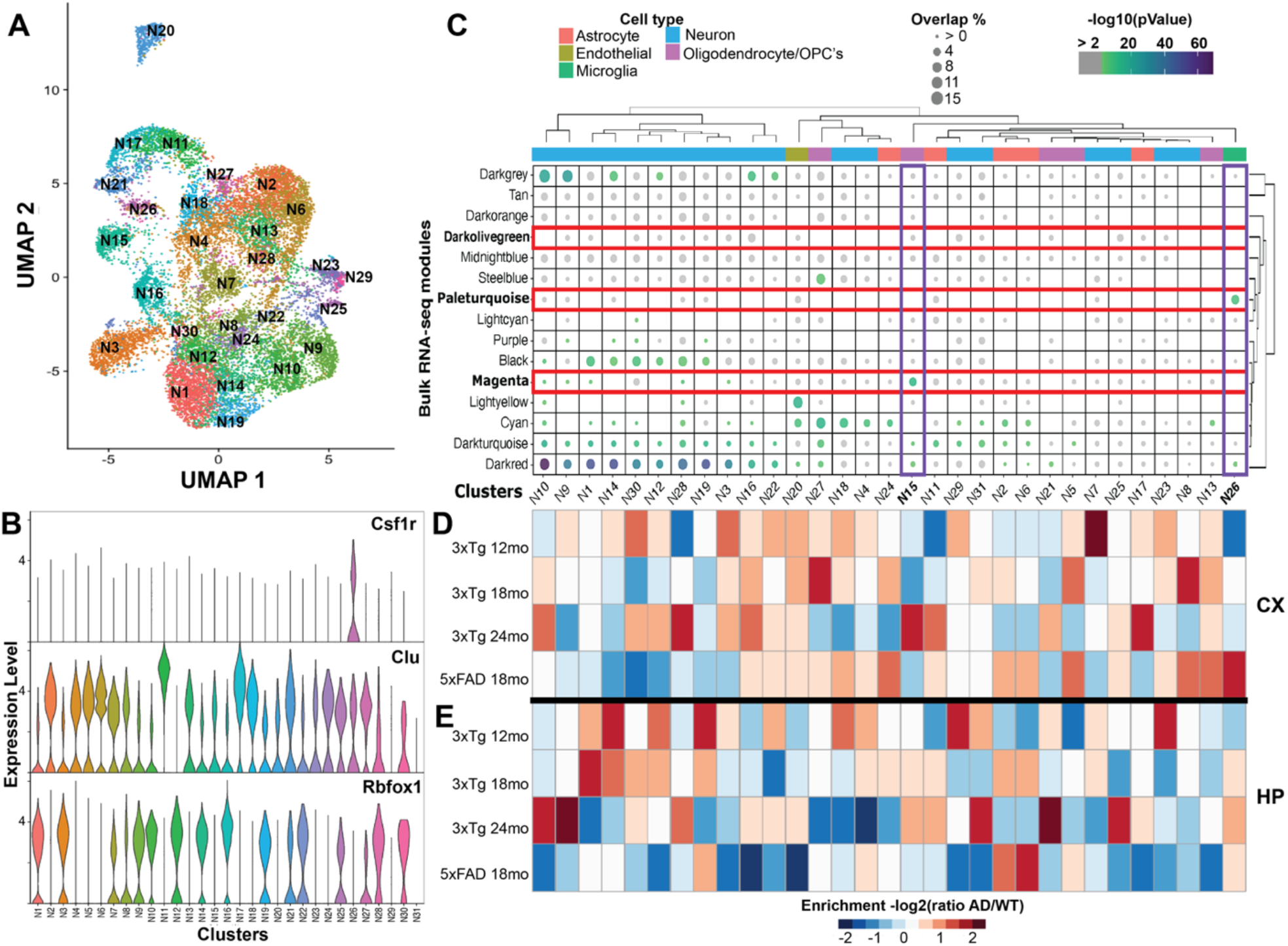
Single-nucleus RNA-seq uncovers astrocytic diversity during neurodegeneration. A) UMAP of 16,482 nuclei colored by clusters. B) Gene expression of markers for microglia (*Csf1r*), astrocytes (*Clu*), and neurons (*Rbfox1*). C) Comparison of cell clusters with 3xTg-AD bulk RNA-seq modules. Sizes of the circles represent percentage of overlap and colors indicate significance of the overlap, where grey is not significant. Enrichment of the genotypes contributing for each cluster in D) cortex and E) hippocampus.

We further compared the genes upregulated in our single-nucleus clusters to our 3xTg-AD WGCNA modules (Figure 2C). Clusters N15 and N26 showed high overlap and significance with the 3xTg-AD magenta (enriched for myelination genes) and paleturquoise modules (enriched for activated microglia genes), confirming the cell identities assigned as oligodendrocytes and microglia, respectively. On the other hand, the darkolivegreen module, which is expressed in young (4 month-old) 3xTg-AD mice, did not show any significant overlap with our snRNA-seq data. The 3xTg-AD darkred module significantly overlapped with eleven neuronal clusters and the black module with 6 other neuronal clusters. Although these modules did not show a particular set of biological functions in the 5xFAD mice (Figure 1G), our results suggest that they might be harboring genes relevant for pathology in the 3xTg-AD.

To identify potential clusters enriched in either 3xTg-AD or 5xFAD mice, we calculated the ratio of “proportion of cells harvested from female AD mouse models” to “proportion of cells harvested from female WT mice” (AD/WT) in each cluster in cortex (Figure 2D) and hippocampus (Figure 2E). When adding male samples, we did not observe significant changes, likely due to the overrepresentation of females in our dataset (Figure S4, Table S3). We observed a substantial increase in microglia enrichment (N26) in 18-month 5xFAD cortex and hippocampus, while in the 3xTg-AD we observed a stronger microglial enrichment later in life, at 24 months. We identified a significant enrichment for astrocyte clusters during neurodegeneration in the 5xFAD specifically in N2 and N6, which mostly originated from the hippocampus (Figure 2E). We observed an increased enrichment for oligodendrocytes (N15) in an age-dependent manner in the hippocampus of 3xTg-AD, but not in the 5xFAD. These results correlate with the increased gene expression patterns of the 3xTg-AD magenta module genes, which relates to myelination. These findings suggest that oligodendrocytes might be important players for the 3xTg-AD pathology.

### In silico trajectory analysis shows a common path of activation between 3xTg-AD and 5xFAD glial cells

We identified substantial expression changes in our astrocyte and oligodendrocyte/OPC snRNA-seq clusters and thus focused on identifying changes in those populations that could be relevant to the progression of AD. To order these cells within a pseudotime, we performed an *in silico* trajectory analysis on astrocyte and oligodendrocyte/OPC clusters using monocle2 ^43^. The resulting trajectory showed the populations of astrocytes at the beginning or root of the pseudotime course, while oligodendrocytes/OPC were placed at the end of the trajectory. Monocle clustering identified at least 5 different potential astrocyte states (Figure 3A-B). Markers relevant to oligodendrocytes, astrocytes, and activated astrocytes were used to calculate their smooth transition across the pseudotime (Figure 3C). Genes *Mobp, Mog, Cspg4*, and *Pdgfra* showed the highest expression at the end of the pseudotime (SG5 and SG6) where the Oligodendrocytes/OPC’s are located. Other genes found in our bulk oligodendrocyte module (magenta) such as *Klk6, Mal, Plp1, Sox10*, and *Myrf* showed increased expression in the oligodendrocyte pseudotime. Neuronal loss has been associated with overexpression of oligodendrocyte markers, such as *Plp1* ^44^. Astrocytic markers *Aqp4* and *Clu* appear at the beginning of the pseudotime, in the SG1 state, which is enriched for 3xTg-AD-derived cells. We observed increased expression of reactive astrocyte genes, such as *Cd9, Lcn2, Vim, Cd44*, and *Cstb* as the cells advance in the pseudotime course, especially in the transitional stages at the end of SG1 and beginning of SG7 (Figure 3A-B). Cells from the 3xTg-AD or 5xFAD samples showed distinct transcriptional profiles during the transitional states. 5xFAD-derived nuclei from astrocytic clusters N2 and N6 were enriched in SG7 and surprisingly showed expression of *Thy1* in similar pattern to the astrocyte activation genes *Serpina3n, Cstb*, and *Serping1* (Figure 3C). Our results show that many transitional astrocytes harvested from 5xFAD mice were already activated, whereas very few 3xTg-AD transitional astrocytes expressed markers of activation. This suggests that 5xFAD-derived astrocytes may activate earlier in response to the aggressive early plaque accumulation seen in the 5xFAD, whereas in the 3xTg-AD mice astrocytes activate later in life coinciding with the slower age-related pathology development in this model.

**Figure 3.**
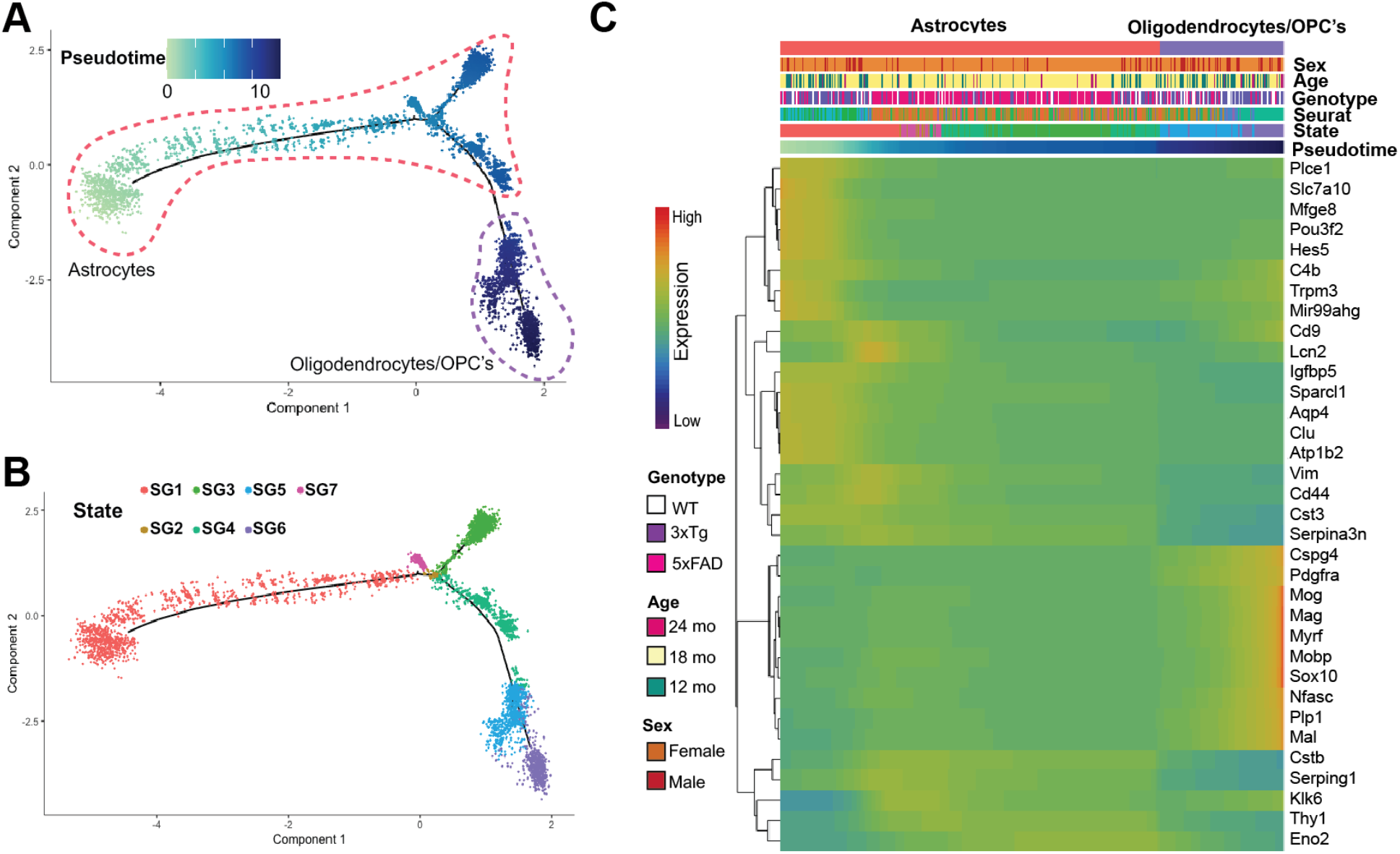
Glial trajectories in AD mouse models display a common path for astrocyte activation. A) Cellular trajectory as ordered by monocle (5,391 cells). B) States identified within the cellular trajectory. C) Expression of marker genes of oligodendrocytes and astrocytes across pseudotime.

### Microglial profiles and trajectories during neurodegeneration

We analyzed the diversity of microglial populations during late stages of neurodegeneration by separately isolating and characterizing microglial cells from the cortex and hippocampus of 12-, 18-, and 24-month 3xTg-AD animals, as well as an 18-month-old 5xFAD female. Single-cell analysis of 23,666 cells identified 23 clusters, which included two non-microglial populations (C18 and C20) representing astrocytes and endothelial cells, respectively (Figure 4). We identified 5 DAM clusters based on high *Cst7* expression and subsets of these clusters also expressed other DAM genes, such as *Itgax* (Figure S5). *Itgax*, also known as *Cd11c*, showed higher expression not only during disease progression, but also during early development. Along with *Clec7a, Itgax* is overexpressed in proliferating microglia that are highly phagocytic during development ^17^. We identified two clusters (C14 and C16) mainly enriched in the 3xTg-AD model that do not express typical DAM genes (Table S4). These two clusters express combinations of *S100a6, S100a9*, and *S100a8*. Genes *S100a9* and *S100a8* have been reported as general microglia markers in human AD data ^31^. Thus, C14 and C16 could potentially represent subpopulations of microglia that are specific to the 3xTg-AD model.

**Figure 4.**
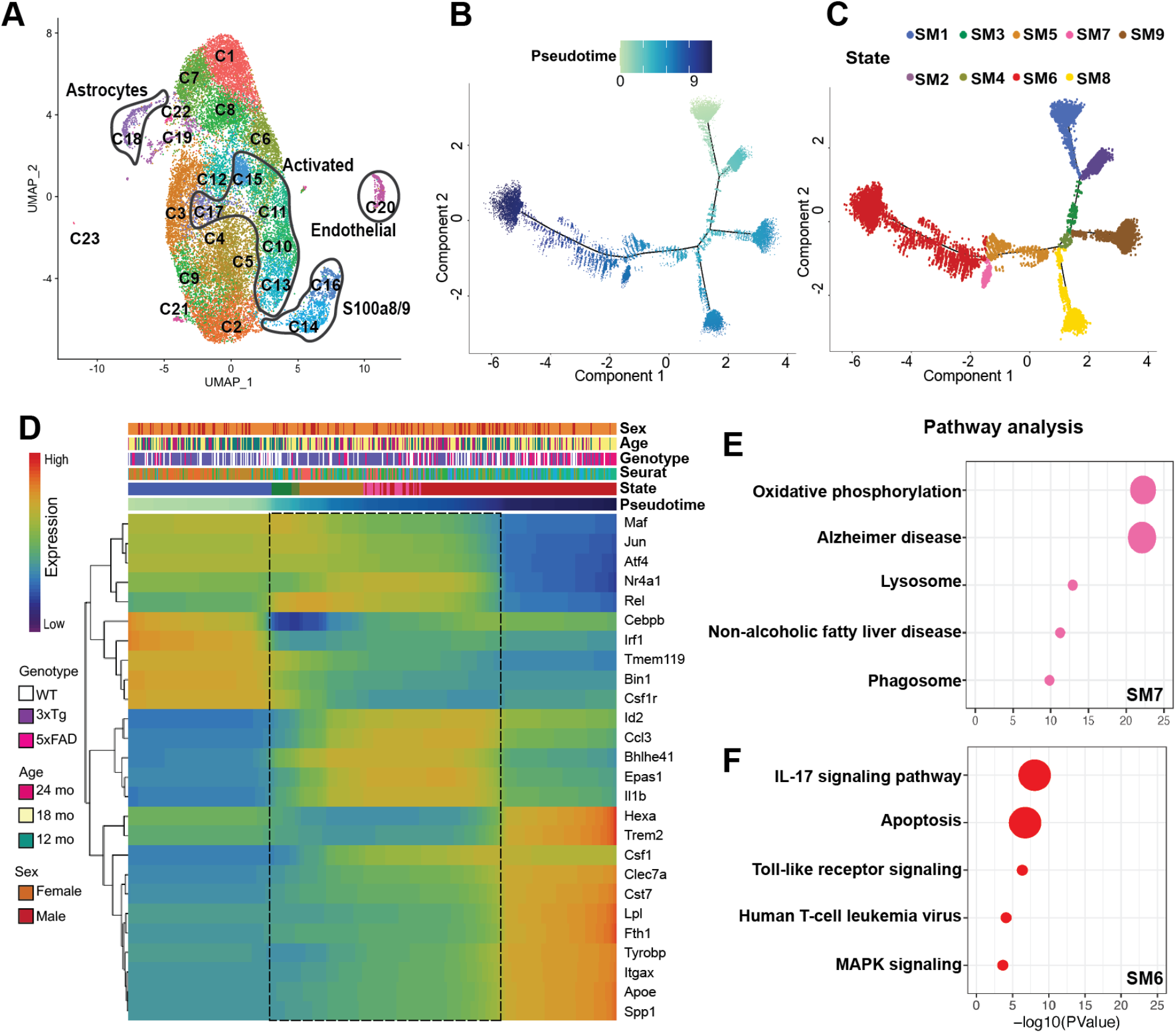
Microglial trajectories during aging. A) UMAP for 23,666 cells across 23 clusters. B) Ordering of microglial cells across pseudotime. C) States identified across the microglial trajectories. D) Smooth expression of genes differentially expressed across pseudo time. Top 5 biological processes associated with the differentially expressed genes in E) SM6 and F) SM7. Intermediate states are delimited by a dashed black box.

DAMs usually overexpress *Cst7, Clec7a, Spp1*, and *Gpnmb* genes. Thus, we sought to identify the cellular trajectories leading to the DAM state and the distinct paths that microglia can take during AD progression. We removed all non-microglial cells and applied Monocle2 to infer trajectories in the 3xTg-AD time course and the 5xFAD timepoint. We found that most microglia branch into four paths (Figure 4B). In addition, we identified nine different states along the pseudotime (Figure 4C). SM6 and SM7 were enriched by the 3xTg-AD and the 5xFAD genotype (Figure S6). To analyze the paths that microglia can take during AD progression, we plotted the expression of markers of homeostatic microglia and DAM across the pseudotime. Genes associated with homeostatic microglia, such *as Tmem119, Csf1r*, and *Irf1* ^45^ showed highest expression at the beginning of the pseudotime. Transcription factors that are involved in microglial functions such as *Cebpb, Maf, and Atf4* ^46^ were highly expressed at the beginning of the trajectory, but their expression decreased along the pseudotime. We observed an increased expression of DAM signature genes at the end of the pseudotime (Figure 4D), which was divided into two states, SM6 and SM7. Interestingly, Csf1r and its ligand *Csf1* showed opposite patterns in the pseudotime course. *Csf1* expression increased along the pseudotime course well-before DAM signature genes increased in expression. *Csf1* is described as a gene involved in homeostasis and promotion of microglial proliferation ^45^. Alternative splicing has been shown to produce two different isoforms of *Csf1*, which could be soluble or insoluble and thus their products are either efficiently or inefficiently released from the cell surface, respectively ^47^. Inclusion of a longer version of exon 6 gives rise to a promptly released growth factor, whereas a shorter version is more slowly processed. Human AD samples show an increase in *Csf1* expression, suggesting an imbalance in this signaling pathways during AD. Another distinctive feature of the early stages seen in our pseudotime course is an increased expression of the *Il1b* gene. We also see an increased expression of multiple TFs, such as *Bhlhe41, Nr4a1*, and *Id2*, whose expression is higher at intermediary states that precede the DAM cells. Interestingly, the expression of these TFs is reduced at the DAM stage. C*ebpb*, which is known to regulate immune genes, including genes critical for glial activation after LPS stimulation ^48^, showed higher expression at the beginning of the trajectory in the 3xTg-AD. This increased expression could indicate a response to subtle neuroinflammation, and the decrease at the end of the trajectory might be result of a negative regulatory loop.

We sought to investigate the distinctions between the two main activated states of microglia (SM6 and SM7) by performing a differential marker analysis. GO terms and pathways enriched in SM6 are mainly associated with interleukin signaling, apoptosis, and toll-like receptors, whereas SM7 is enriched for oxidative phosphorylation, phagosome, and lysosome (Figure 4E-F). Overall, we observed homeostatic and activated clusters that are part of a main trajectory towards neurodegeneration. The 5xFAD-derived microglia showed the strongest DAM signature profile, while the 3xTg-AD-derived microglia was mainly present in intermediate states on their way towards neurodegeneration. SM6 and SM7 states are involved in cytokine stimuli and oxidative phosphorylation, respectively (Figure 4E-F). SM6 showed the strongest activation and thus the main path towards neurodegeneration. These findings suggest a slower pace of microglia activation in the 3xTg-AD when compared to the 5xFAD, in concordance with the stronger and earlier pathology seen in the latter.

### scATAC-seq reveals DAM-specific regions of accessible chromatin in the cortex of a 24-month-old female 3xTg-AD mouse

We performed scATAC-seq in a half cortex of a 24-month-old 3xTg-AD mouse to investigate changes in chromatin accessibility in activated microglia. We recovered 4,451 cells that were distributed into 10 distinct clusters using Signac (Figure 5A). We identified open chromatin regions driving the formation of each cluster (Figure 5B). We further mapped scATAC clusters to their equivalent scRNA-seq clusters using the Seurat anchor finding function (Figure 5C). Most of the clusters found in the scRNA-seq dataset were distributed across 2 main branches in the UMAP. RNA cluster C14 is the only cluster that has a clear correspondence to scATAC-seq cluster A8 (Figures 5A, C). Interestingly, all of our activated microglial clusters from scRNA-seq mapped into A2. This suggests that although we found distinct transcriptional activation states, the chromatin profile is mostly conserved among them. We performed GO analysis on open chromatin regions specific for A2 using GREAT ^49^, which recovered significant enrichment in genes related to response to external stimulus, LPS, and defense response (Figure 5D). An enrichment for LPS response/inflammation suggests the opening of chromatin regions near genes associated with microglial activation. A regulatory element located 18 kb upstream of *Csf1* is associated with A2. Predicted expression levels of *Csf1* are higher for A2, matching the observed pattern in our scRNA-seq where activated microglia, present at the end of our pseudotime course, show an increase in *Csf1* (Figure 5E). We conclude that 3xTg-AD-derived activated microglia have a distinct open chromatin profile compared to homeostatic microglia.

**Figure 5.**
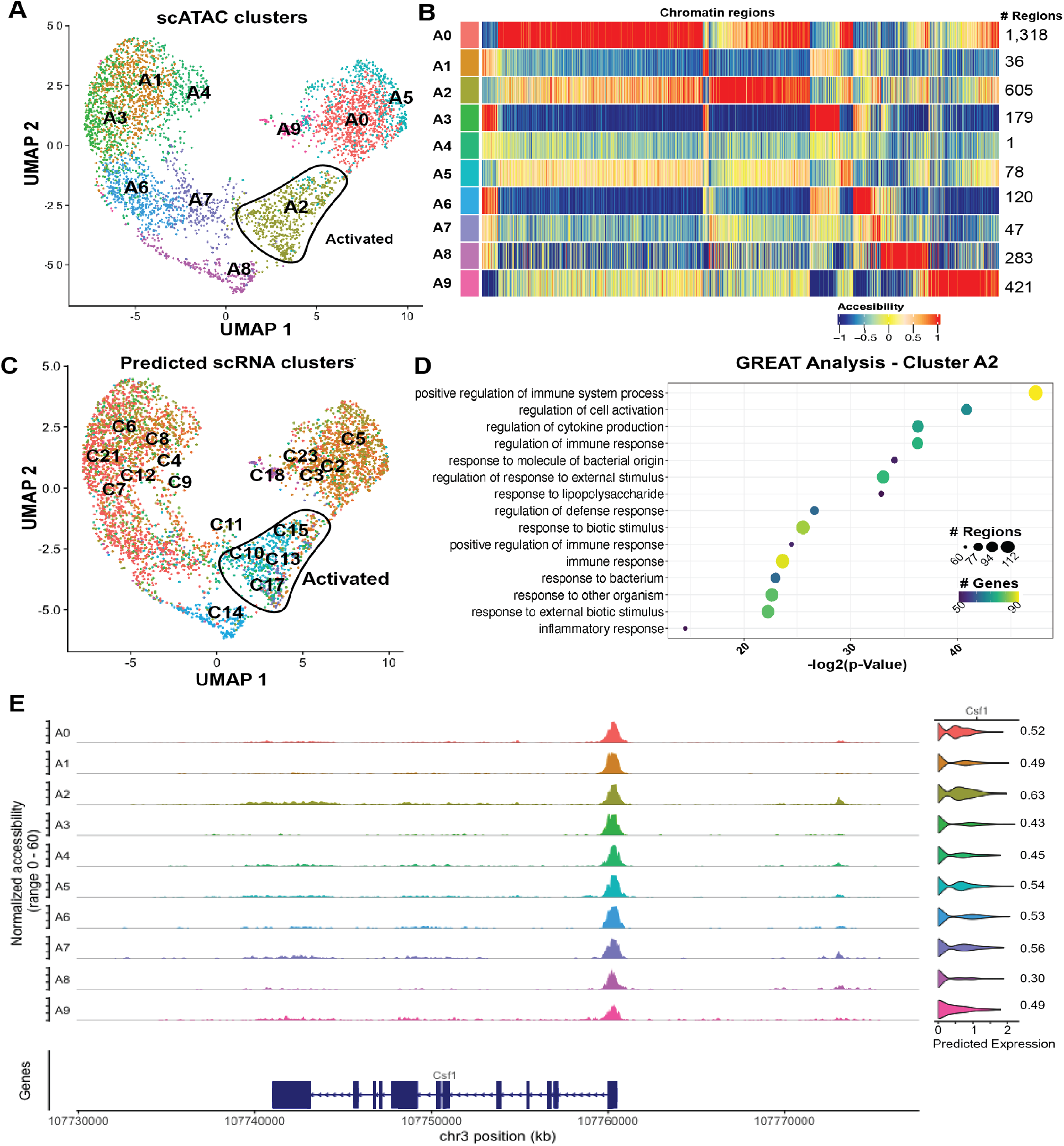
Chromatin accessibility of activated microglia derived from 24-month-old 3xTg-AD mouse. A) UMAP highlighting clustering of 4,451 microglial chromatin profiles. Activated microglia are circled in black and labeled. B) Chromatin accessible regions per cluster. C) UMAP labeled by predicted labels from scRNA-seq microglial data (Figure 2A). Activated microglia are circled in black and labeled. D) GO Term enrichment for cluster A2 specific chromatin regions. Light color represents higher number of genes. D) *Csf1* coverage plot in ATAC-seq clusters. Violin plots represent the predicted expression. Red box highlights candidate regulatory regions potentially contributing to predicted gene expression activity.

## DISCUSSION

In this work, we describe the progression of AD-like pathology through the 3xTg-AD mice lifespan using multiple functional genomics approaches. Transcriptomic profiles at the bulk level in the hippocampus of 3xTg-AD female mice showed an overlap with gene modules previously described in humans and the 5xFAD model. The importance of glial cells is confirmed as genes related to myelination and immune response are conserved between species and/or models. One of the major characteristics of AD is the loss of myelination ^50^, yet the oligodendrocyte module (magenta) shows an increase over time in the 3xTg-AD mice. The increase in gene expression of oligodendrocyte-associated genes does not imply an increase in cell numbers but suggests that oligodendrocytes could be attempting to remyelinate neurons with 3xTg-AD aging. Some of the genes present in the oligodendrocyte (magenta), neuron (darkolivegreen), and glial (paleturquoise) modules overlap with neuronal system modules described in AD samples, suggesting that changes in neuronal function could account for the behavioral phenotypes of the 3xTg-AD mice.

Genes associated with oligodendrocytes were enriched in the 3xTg-AD hippocampus and their expression increased with age, which was not seen in the 5xFAD mice. A decrease in the oligodendrocyte population has been reported in the aging 3xTg-AD mice ^26^, whereas an increase in OPCs was observed in the APP/PS1 model ^50^. The reason for this discrepancy in oligodendrocyte response in AD models is still unknown, but it could be associated with when and how fast the pathology develops in different models, or with the development of hyperphosphorylated tau and neuronal damage. Expression of inflammatory genes, as well as activated microglia-, activated astrocyte- and oligodendrocyte-associated genes increase with aging in the hippocampus of the 3xTg-AD model. Hence, microglia, astrocytes, and oligodendrocytes seemed to respond to injury and might play important roles in the pathology of the 3xTg-AD. These results can contribute to elucidating the role of oligodendrocytes in AD pathology.

5xFAD hippocampus-derived astrocytes from clusters N2 and N6 showed high Thy1 expression, which was also detected in a few 3xTg-AD astrocytes. *Thy1* is usually expressed by neurons only, but some studies have also detected *Thy1* expression in glial cells ^51^. Transgenes from both 5xFAD and 3xTg-AD models are regulated by the mouse *Thy1* mini gene ^52–54^. The overexpression of *Thy1* in our data could be due to greater expression of the transgene in both models. Astrocytes expressing *Thy1* were substantially enriched in the 5xFAD mice, which are known to have increased number of astrocytes ^6^, compared to the 3xTg-AD. Thus, *Thy1* might serve as an additional marker of astrocyte activation.

3xTg-AD and 5xFAD mice seem to share some inflammatory signaling pathways and microglia harvested from hippocampus of both models shared a common trajectory of activation. However, with 5xFAD-derived microglia seem to be activated earlier and also show stronger expression of DAM genes versus 3xTg-AD-derived microglia, which showed slower and weaker activation. These findings agree with the presence of much earlier pathology in the 5xFAD mice. We also observed an interesting dynamic between Csf1r and its ligand *Csf1*. While Csf1r is highly expressed in homeostatic conditions, *Csf1* is expressed in activated microglia in our data. Decreased expression of *Csf1r* has been previously reported in AD, whereas an increase in *Cfs2*, which is also a *Csf1r* ligand, has been described as crucial for inflammation and neurodegenerative diseases ^55^. The polemic of macrophage activation can extend to our data in the context of microglia and their different reactions to injury. While microglia activation has a clear signature (DAM), the microglial responses to different combination of pathologies, such as plaques and/or tangles might be just a continuum of the DAM activation patterns.

The addition of chromatin data from a 24-month-old 3xTg-AD female cortex provides an additional layer of information to our data. While we only have one time point and we do not have a matching WT, we were still able to identify a specific chromatin profile for most of the activated microglial clusters. Multiple clusters identified as activated microglia in the scRNA-seq dataset are correlated with the snATAC-seq A2 cluster, implying that the diverse activated responses share a similar chromatin profile that is distinct from homeostatic microglia. Association of ATAC-seq data from human AD samples with targets obtained from GWAS studies showed that some of the relevant AD SNPs are present in open chromatin regions of glial cells ^56^. The full characterization of open chromatin regions can help identify additional AD risk factors. Here, we provide characterization of the transcriptomic profiles of microglia, astrocytes, and oligodendrocytes during disease progression in the 3xTg-AD mouse model. We also provide insight into the chromatin profiles of activated microglia in late stage of neurodegeneration in the 3xTg-AD mouse. Our results provide a valuable resource to help elucidate the progression of neurodegeneration in the presence of plaques and tangles and can potentially improve translatability.

## METHODS

### Animals

All animal experiments were approved by the UC Irvine Institutional Animal Care and Use Committee and were conducted in compliance with all relevant ethical regulations for animal testing and research. 3xTg-AD mice (B6;129-Psen1tm1MpmTg(APPSwe,tauP301L)1Lfa/ Mmjax, JAX MMRRC Stock# 034830) and their WT littermates were bred, housed, and taken care by the animal facility at UCI. For bulk RNA-seq the animals were sacrificed via CO2 inhalation and transcardially perfused with 1X phosphate buffered saline (PBS). For single-cell experiments, animals were sacrificed and perfused as previously described ^6^. Half of the brain was kept in a Hank’s balanced salt solution (HBSS) without Calcium and Magnesium and used fresh for single-cell experiments while the other half was snap frozen and stored for single-nucleus isolation.

### Bulk RNA sequencing

Bulk RNA-seq libraries were prepared as described previously ^6^. Briefly, the libraries were constructed following the Smart-seq2 protocol ^57^ utilizing the Illumina Nextera DNA Sample Preparation kit. The quality of the libraries was assessed using the Agilent 2100 Bioanalyzer and the libraries were normalized based on Illumina KAPA Library Quantification Kit results. Final libraries were sequenced using the NextSeq 500 (Illumina). Samples’ sequences were aligned to the mouse reference genome mm10 using STAR (2.5.1b-static) ^58^ and gene expression was quantified using RSEM (1.2.22) ^59^.

### WGCNA analysis

A gene expression matrix (created by RSEM) and filtered by genes with more than 1 TPM was used as an input of weighted gene correlation network analysis (WGCNA)^60^. After removing outlier samples, power 14 was chosen to reach a scale free network. We detected 15 modules with a set of genes with similar expression patterns. We calculated the Pearson correlation for genes in each module followed by calculation of Fisher’s asymptotic p-value for given correlations. The 3xTg-AD modules with significant correlations (p-value < 0.05) with genotype (magenta, darkolivegreen and paleturquoise) were plotted using an eigengene profile. The analysis outputs a heatmap in which each column represents the samples, and each row represents the genes and eigengene barplot. EnrichR ^61^ was used for GO term analysis in significant 3xTg-AD modules (magenta, darkolivegreen and paleturquoise) with adjusted p-value < 0.05.

### Comparison of 3xTg-AD modules to previously identified human AMP-AD and 5xFAD modules

The modules obtained from WGCNA analysis in 3xTg-AD samples were compared to previously identified 5xFAD modules ^6^ and to human AMP-AD modules ^42^ by calculating genes overlap and significance (p-value) as previously described ^6^. Briefly, the gene overlap was calculated by the formula (| A ∩ B|) / |A| * 100, in which A=Genes present in a 3xTg-AD module and B=genes present in a single-cell cluster. For significance we used Fisher exact test ([N - |A∪B|, A-B; B-A, |A∩B|], in which N = number of all genes, A = gene set in each 3xTg-AD gene lists and B = gene set in each AMP-AD module, or B= gene set in each 5xFAD module.

### Single-nucleus RNA-seq

Half cortex and half hippocampus were snap frozen before nuclei extraction. Nuclei were extracted as described in the “Nuclei Extraction” section of SPLiT-seq Protocol V3.0 ^62^. Buffer volumes were adjusted according to the tissue lysed. Homogenates were filtered with 30μm MACS SmartStrainers (Cat# 130-098-458). The nuclei were resuspended, counted (using the BioRad TC20™ Automated Cell Counter), and diluted in 0.05% Bovine Serum Albumin (BSA) in Phosphate-buffered saline (PBS) to reach 10,000 nuclei per microliter. Each single-nucleus was lysed and barcoded inside the nanodroplets generated by the ddSEQ Single-Cell Isolator. cDNA synthesis, tagmentation and final library generation were performed using the Illumina Bio-Rad SureCell WTA 3’ Library Prep Kit (Cat# 20014280). cDNA concentration and size distribution were assessed using the Agilent 2100 Bioanalyzer. The amount of transposase and buffer (tagmentation mix) were adjusted per sample based on its size measured by Agilent bioanalyzer. The resulting libraries’ size was 900 to 1100 base pair. Single-nucleus libraries were sequenced using a 150-cycle NextSeq 500/550 High Output Kit v2.5 (Cat# 20024907) on the Illumina NextSeq500 platform. The sample’s loading concentration was between 2.0 to 2.4pM and we utilized a custom primer provided with the SureCell WTA 3’ to perform the sequencing run. Each library had a minimum depth of 50 million reads.

### Microglia single-cell RNA-seq

Perfused samples (half cortex or half hippocampus) were dissociated while fresh using Miltenyi Adult Brain dissociation kit (Cat# 130-107-677) and gentleMACS™ Octo Dissociator with Heaters. The dissociated tissues were filtered with 70μm MACS SmartStrainers (Cat# 130-110-916). We proceeded with removal of debris and myelin from the single-cell suspension using myelin beads II (130-096-733). The resulting cells were enriched for microglia with magnetic labeling using CD11b MicroBeads (Cat# 130-093-634). Labeled cells were resuspended in 10-15μl of buffer (0.05% BSA in DPBS) for half hippocampus samples and 20-30μl of buffer for half cortex samples to reach a minimum of 1,000 cells per microliter up to 7,000 cells. After counting the cells and assessing cellular viability using the TC20™ Automated Cell Counter, we profiled the transcriptomes of microglia (single-cell). The following library preparation steps are the same as previously described in the “Single-nucleus RNA-seq” section.

### scRNA-seq and snRNA-seq pre-processing and analyses

Fastq files were processed using the KB_python pipeline, which is based on kallisto bustools ^63^ using the parameters: kb count -x SURECELL --h5ad -t 20. Cells with more than 200 genes and less than 10% mitochondrial reads were utilized in downstream analysis. Samples with more than 100 cells were used for clustering and further analysis. Seurat V4 ^64^ was used to perform normalization, regression, clustering (using the Leiden algorithm), and differential expression analysis of single-cell data. For doublets removal we used the package scD ^65^.

Ratio comparisons were done calculating the proportion in which each genotype/age contributes to each cluster. The proportion of AD genotype was then divided by the proportion of their corresponding WT.

### scATAC-seq experiment

Microglial cells were isolated from half cortex of a 24-month-old 3xTg-AD female mouse, as described in the “Microglia single-cell RNA-seq” section. Cells were collected at the bottom of an Eppendorf tube, all supernatant was removed, and they were snap frozen to be stored at -80 until use. The cells were then resuspended into PBS + 0.1% BSA buffer. The Omni-ATAC version of the Bio-Rad ddSEQ SureCell ATAC-seq (Illumina) workflow was followed ^66,67^, Briefly, the cells were lysed with ATAC-Lysis buffer containing digitonin. The nuclei extracted were washed with ATAC-Tween buffer. To confirm that the cells were lysed we took an aliquot and stained them with Trypan Blue (Bio-Rad) to visualize under the microscope. Around 60,000 nuclei were tagmented. After tagmentation, we amplified the fragments by loading the tagmented sample into the ddSEQ where barcoded beads are added to each nucleus. Leftover tagmented nuclei that were not loaded in the first round of barcoding were frozen and stored for a second replicate. Libraries were assessed in a bioanalyzer to measure concentration and size distribution before being loaded onto the NextSeq500 (Illumina) and sequenced at 1.5 pM loading concentration using a custom primer.

### scATAC-seq processing

Raw fastqs were processed using the Biorad scATAC pipeline. A custom script was used to generate count matrices that were processed with Sinto (https://github.com/timoast/sinto) to generate fragment files used as input for Signac ^64^. After requiring a minimum of 1 cell to present the chromatin regions we used the parameters as follows: nCount_peaks > 3000 & nCount_peaks < 100000 & nucleosome_signal < 4 & TSS.enrichment > 2. We then ran standard processing steps that include *RunTFIDF, RunSVD* and dimension reduction by ‘*lsi*’.

## SUPPLEMENTARY FIGURE LEGENDS

**Figure S1 Correlation matrix per sample used in the WGCNA analysis**. Each row and column represent each one of the samples used for the analysis. Age and genotype are denoted by the colored bars. The highest positive Pearson correlation between samples is represented in bright red.

**Figure S2 Module trait correlation including male samples from hippocampus and cortex**. Rows represent the different gene modules and columns represent different traits. Correlation values are presented by color and the first numerical value in each box. Significance values are represented inside of parentheses.

**Figure S3 Enrichment analysis of the darkolivegreen 3xTg-AD module.** GO terms for the darkolivegreen module. The significance is shown as -log2(p-value). The size of the circle represents the number of genes in the GO Term and color represents the false discovery rate.

**Figure S4 Ratio enrichment in cortex and hippocampus of female and male 3xTg-AD mice in relative to WT**. Each column represents a genotype and age. Rows represent single-nuclei clusters. Positive enrichment for the AD genotype is in red and for WT is in blue.

**Figure S5 Homeostatic microglia and activated microglia markers in scRNA-seq clusters**. Gene expression for homeostatic microglia (*Csf1r*) and activated microglia (*Cst7* and *Itgax)*.

**Figure S6 Genotype enrichment plot in microglial states**. States as described in Figure 2.8. Enrichment of AD/WT is plotted as log2(Ratio) per timepoint and state.

## SUPPLEMENTARY TABLES

***Table S1*** Animals used per experiment for single-cell and single-nucleus

***Table S2*** Top 30 marker genes per cell type in the snRNA-seq data

***Table S3*** Nuclei number per animal and cluster

***Table S4*** Top 10 marker genes per cluster in the scRNA-seq data

## Supplementary figures

**Figure S1.**
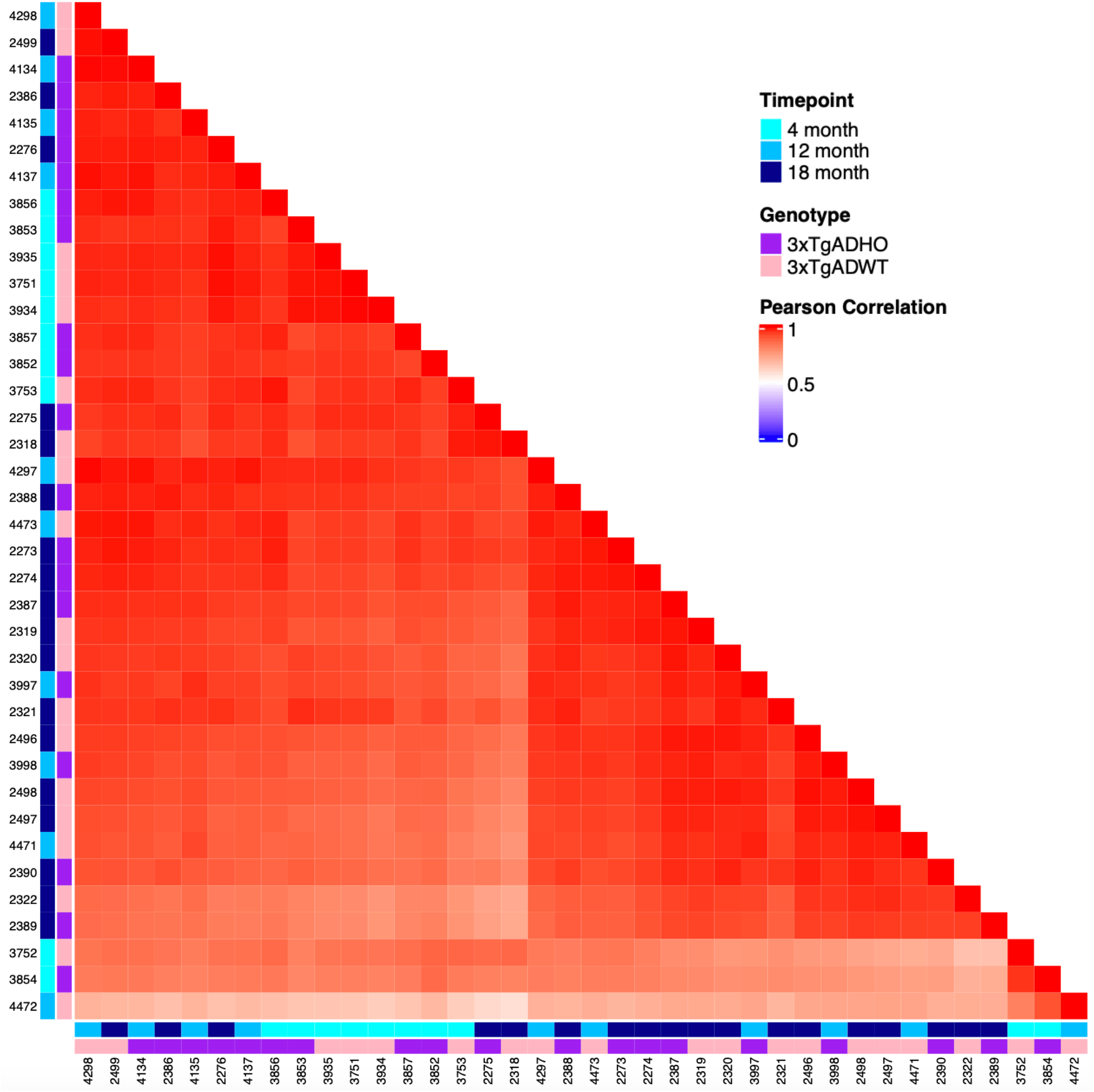
Correlation matrix per sample used in the WGCNA analysis. Each row and column represent each one of the samples used for the analysis. Age and genotype are denoted by the colored bars. The highest positive Pearson correlation between samples is represented in bright red.

**Figure S2.**
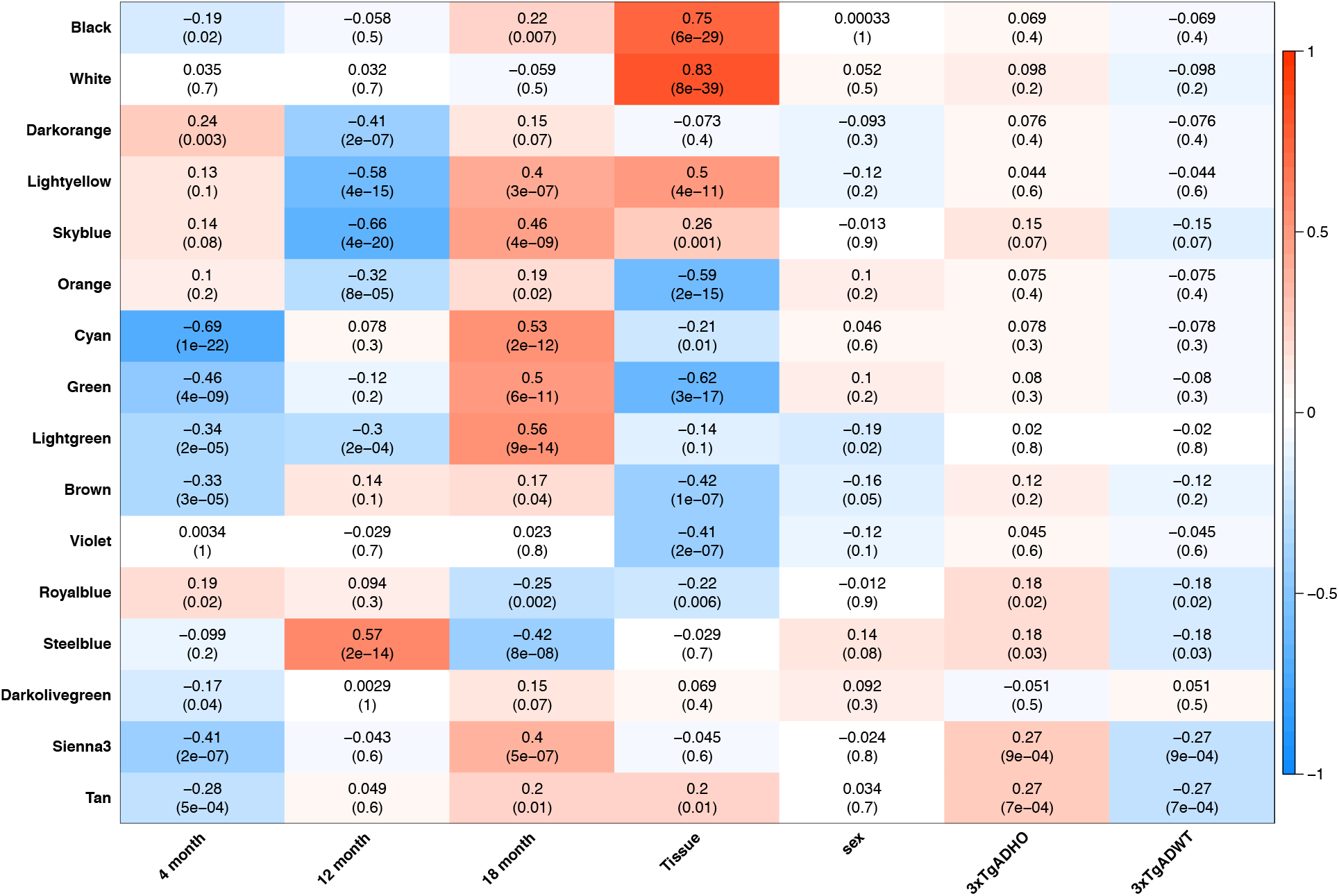
Module trait correlation including male samples from hippocampus and cortex. Rows represent the different gene modules and columns represent different traits. Correlation values are presented by color and the first numerical value in each box. Significance values are represented inside of parenthesis.

**Figure S3.**
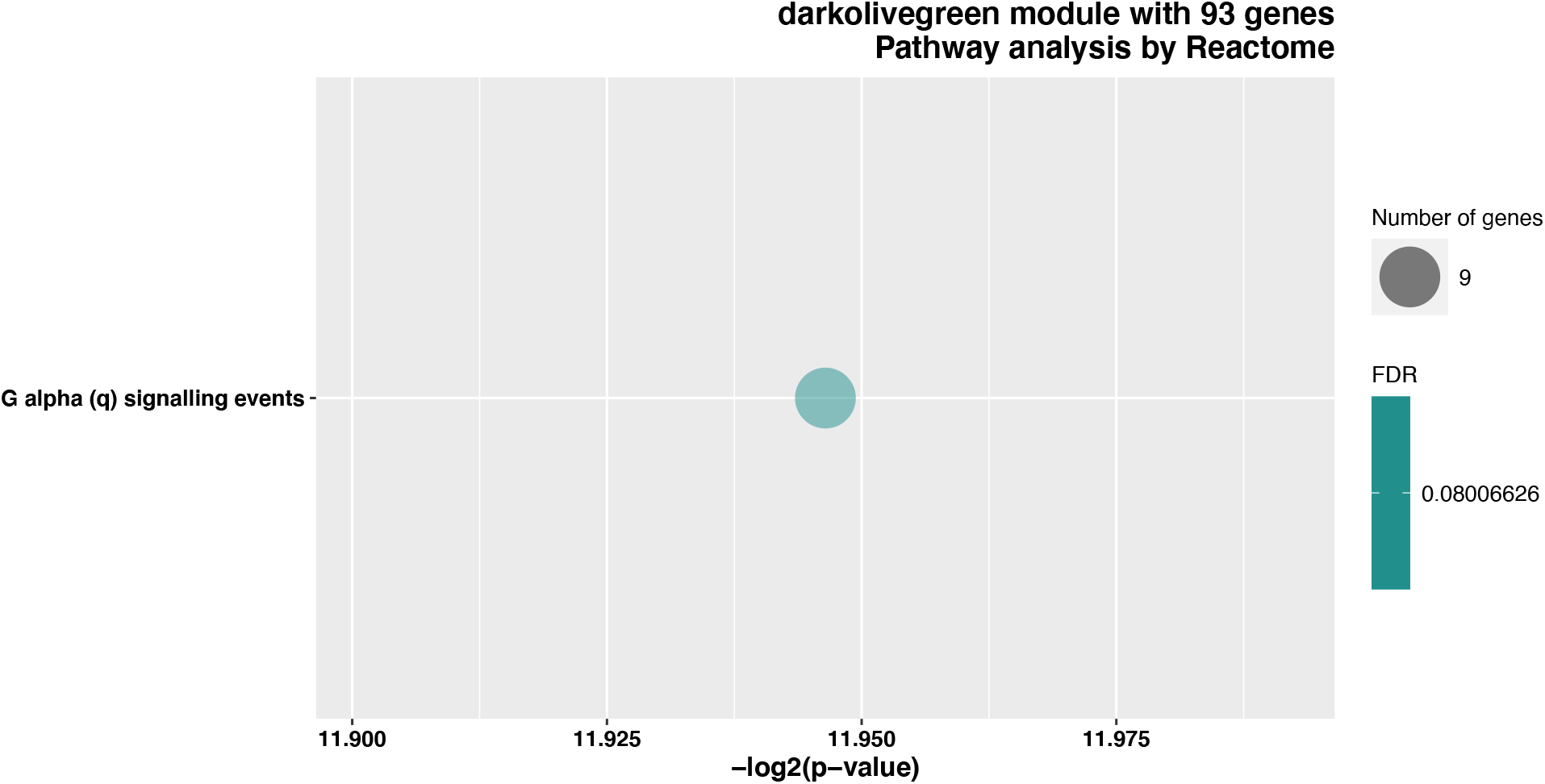
Enrichment analysis of the darkolivegreen 3xTg module. GO term for the darkolivegreen module. The significance in -log2(p-value). Size of the circle represents number of genes in the GO Term and color represents the false discovery rate.

**Figure S4.**
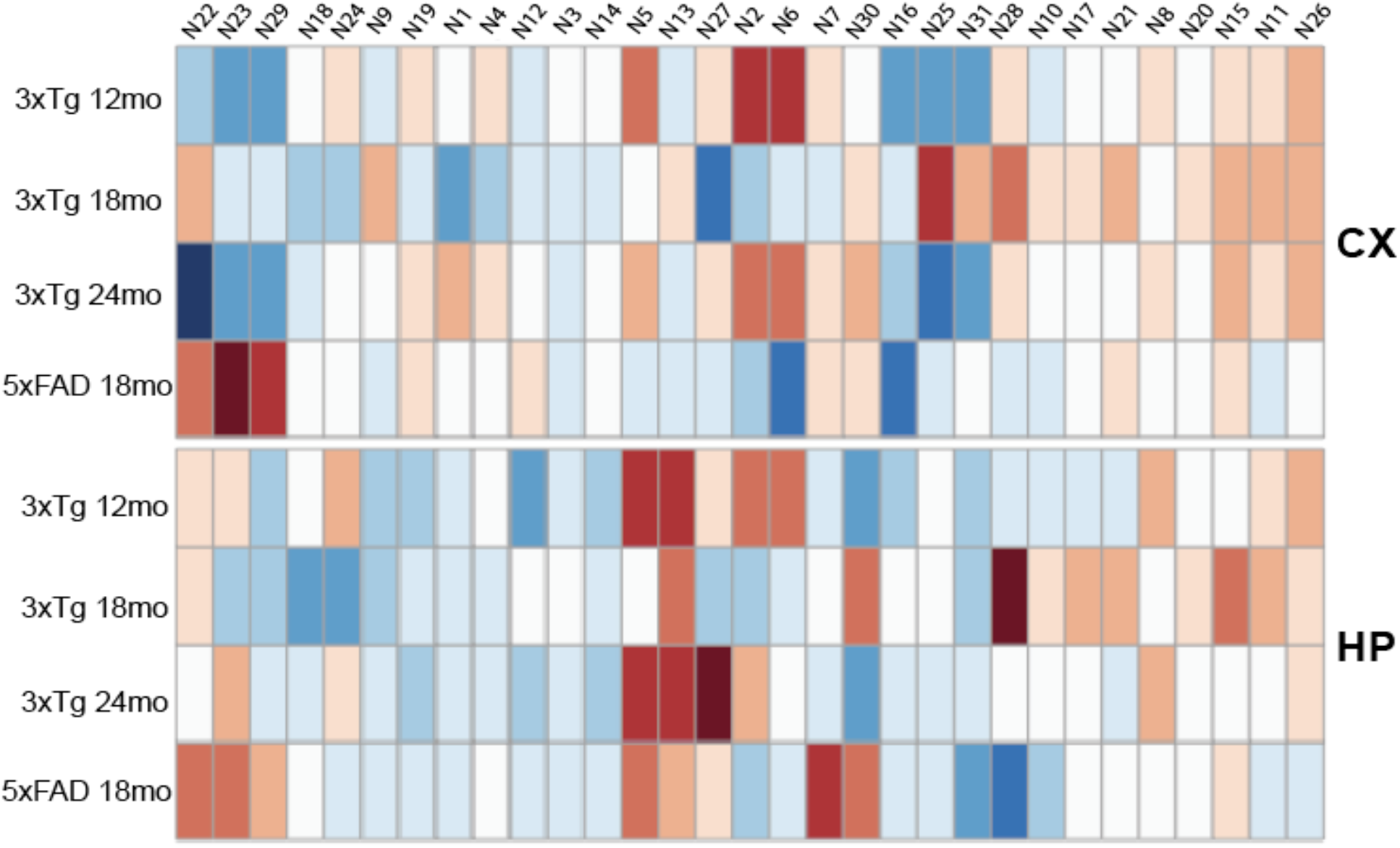
Ratio enrichment in cortex and hippocampus of female and male 3xTg mice in relative to WT. Each column represents a genotype and age. Rows represent single-nuclei clusters. Positive enrichment for the AD genotype is in red and for WT in blue.

**Figure S5.**
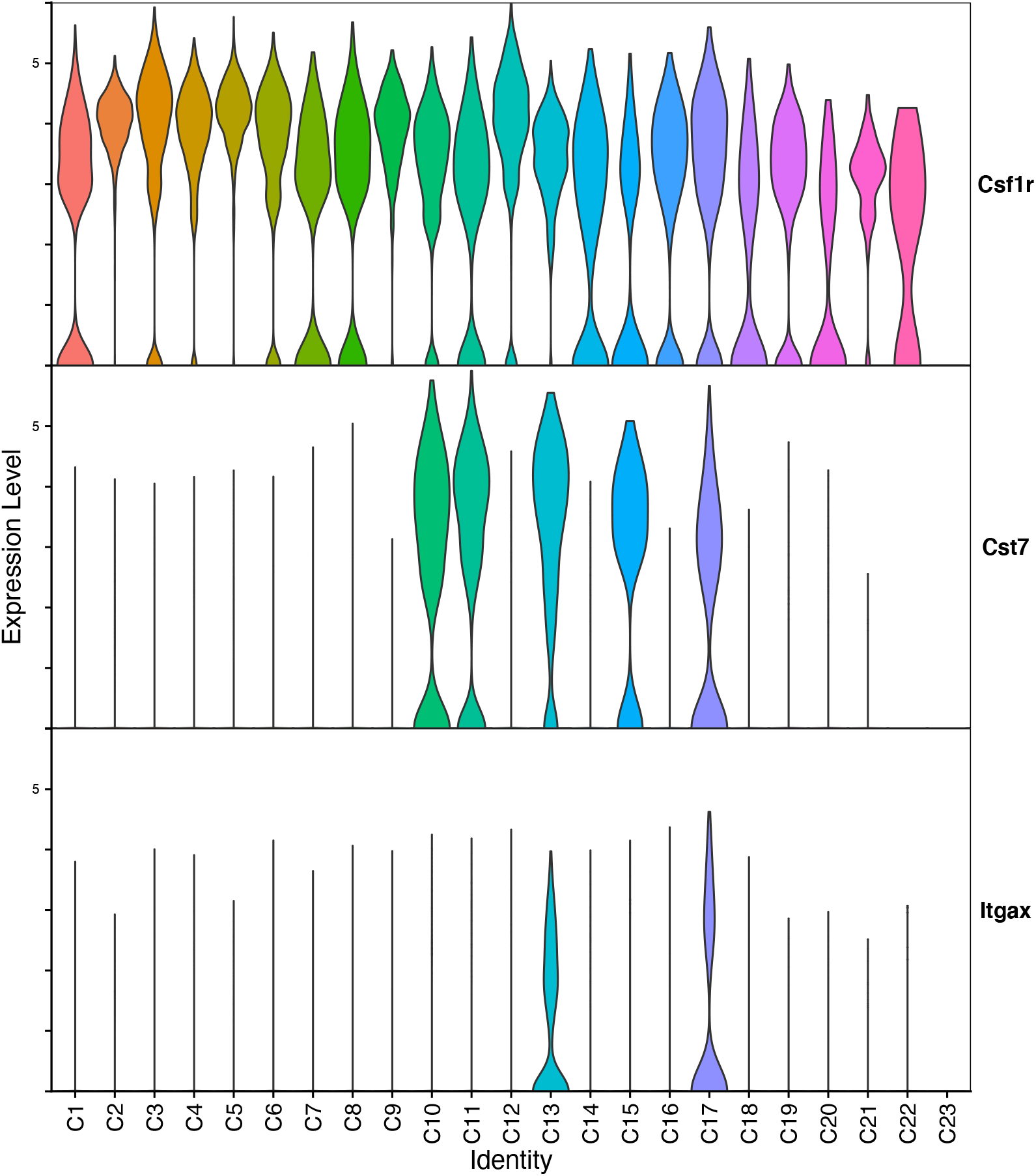
Microglia and activation subtype markers in scRNA-seq clusters. Gene expression for microglia (*Csf1r*) and activation (*Cst7* and *Itgax)*.

**Figure S6.**
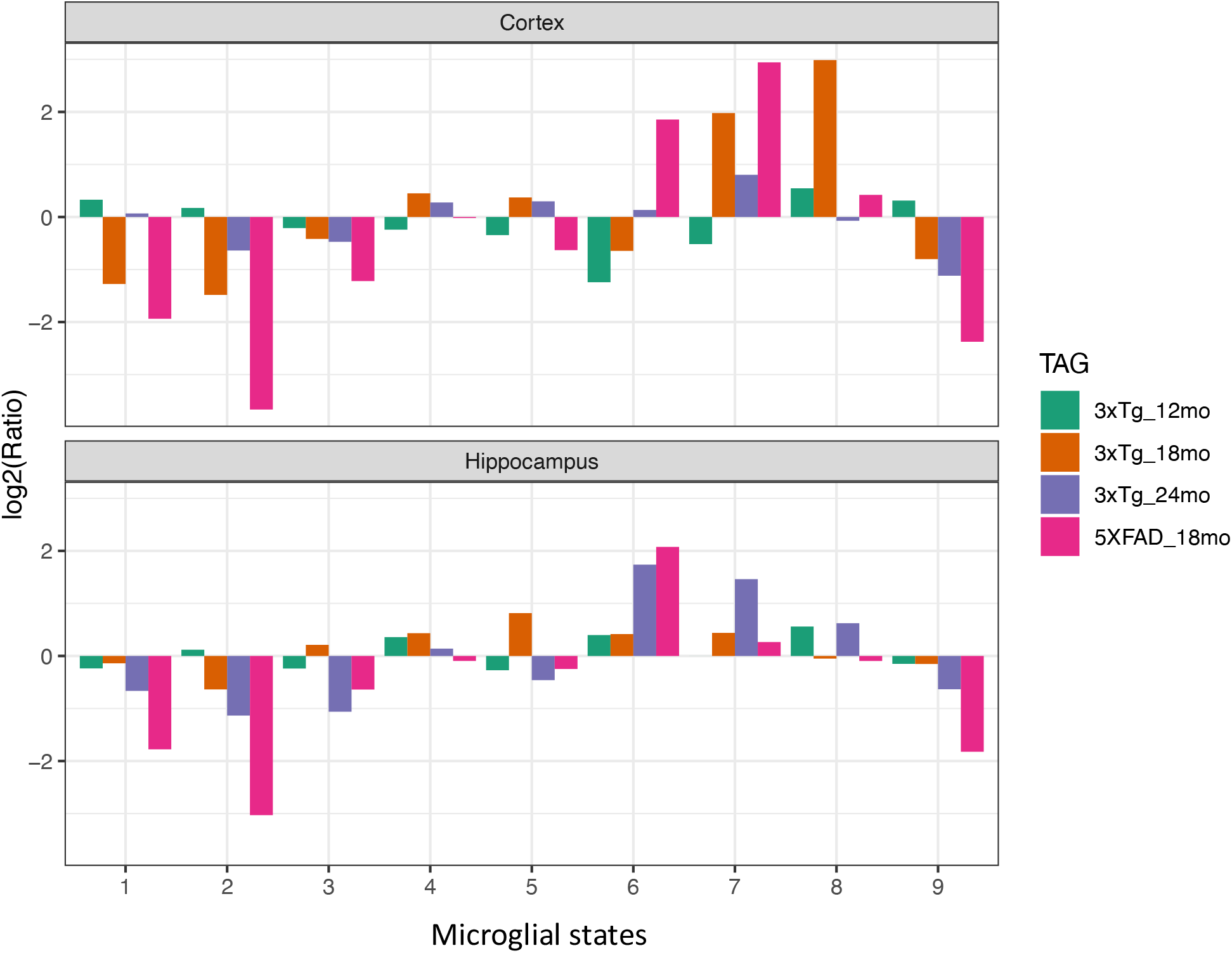
Genotype enrichment plot in microglial states. States as described in Figure 2.8. Enrichment of AD/WT is plotted as log2(Ratio) per timepoint and state.

## Supplementary tables

**Table S1.**
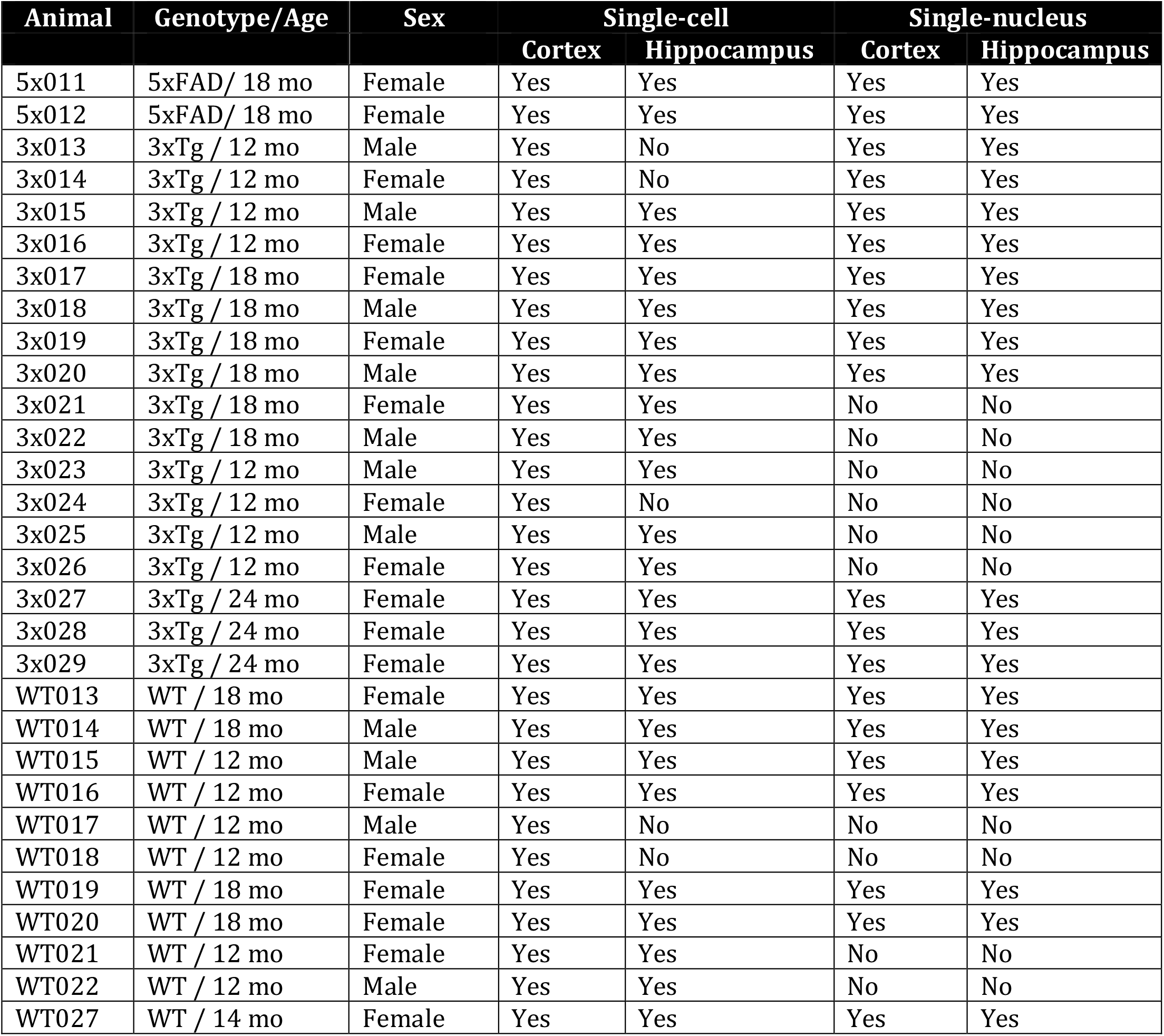
Animals used per experiment for single-cell and single-nucleus.

| Animal | Genotype/Age | Sex | Single-cell |  | Single-nucleus |  |
| --- | --- | --- | --- | --- | --- | --- |
|  |  |  | Cortex | Hippocampus | Cortex | Hippocampus |
| 5x011 | 5xFAD/ 18 mo | Female | Yes | Yes | Yes | Yes |
| 5x012 | 5xFAD/ 18 mo | Female | Yes | Yes | Yes | Yes |
| 3x013 | 3xTg / 12 mo | Male | Yes | No | Yes | Yes |
| 3x014 | 3xTg / 12 mo | Female | Yes | No | Yes | Yes |
| 3x015 | 3xTg / 12 mo | Male | Yes | Yes | Yes | Yes |
| 3x016 | 3xTg / 12 mo | Female | Yes | Yes | Yes | Yes |
| 3x017 | 3xTg / 18 mo | Female | Yes | Yes | Yes | Yes |
| 3x018 | 3xTg / 18 mo | Male | Yes | Yes | Yes | Yes |
| 3x019 | 3xTg / 18 mo | Female | Yes | Yes | Yes | Yes |
| 3x020 | 3xTg / 18 mo | Male | Yes | Yes | Yes | Yes |
| 3x021 | 3xTg / 18 mo | Female | Yes | Yes | No | No |
| 3x022 | 3xTg / 18 mo | Male | Yes | Yes | No | No |
| 3x023 | 3xTg / 12 mo | Male | Yes | Yes | No | No |
| 3x024 | 3xTg / 12 mo | Female | Yes | No | No | No |
| 3x025 | 3xTg / 12 mo | Male | Yes | Yes | No | No |
| 3x026 | 3xTg / 12 mo | Female | Yes | Yes | No | No |
| 3x027 | 3xTg / 24 mo | Female | Yes | Yes | Yes | Yes |
| 3x028 | 3xTg / 24 mo | Female | Yes | Yes | Yes | Yes |
| 3x029 | 3xTg / 24 mo | Female | Yes | Yes | Yes | Yes |
| WT013 | WT / 18 mo | Female | Yes | Yes | Yes | Yes |
| WT014 | WT / 18 mo | Male | Yes | Yes | Yes | Yes |
| WT015 | WT / 12 mo | Male | Yes | Yes | Yes | Yes |
| WT016 | WT / 12 mo | Female | Yes | Yes | Yes | Yes |
| WT017 | WT / 12 mo | Male | Yes | No | No | No |
| WT018 | WT / 12 mo | Female | Yes | No | No | No |
| WT019 | WT / 18 mo | Female | Yes | Yes | Yes | Yes |
| WT020 | WT / 18 mo | Female | Yes | Yes | Yes | Yes |
| WT021 | WT / 12 mo | Female | Yes | Yes | No | No |
| WT022 | WT / 12 mo | Male | Yes | Yes | No | No |
| WT027 | WT / 14 mo | Female | Yes | Yes | Yes | Yes |

**Table S2.**
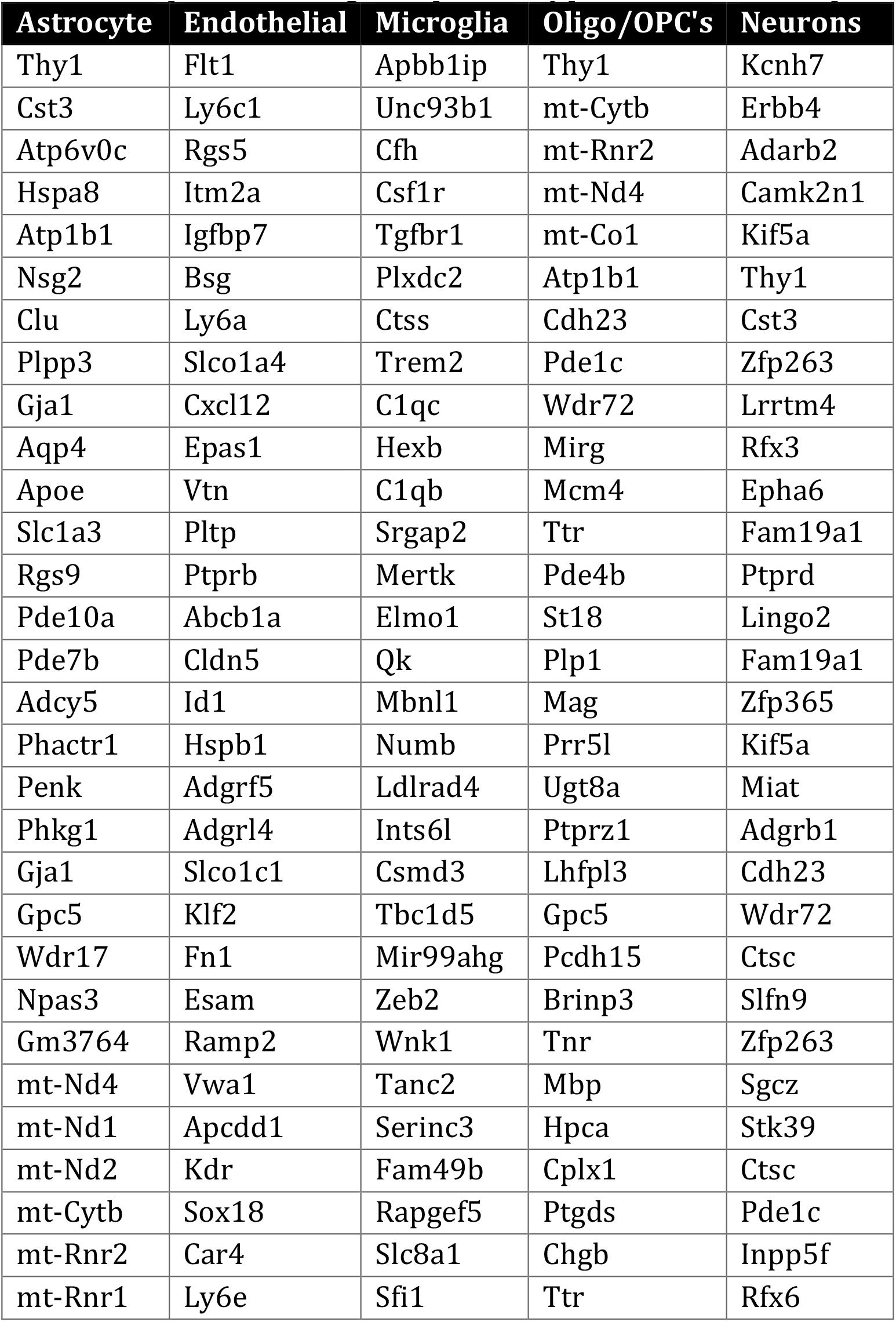
Top 30 marker genes per cell type in the snRNA-seq data.

**Table S4.** Top 10 marker genes per cluster in the scRNA-seq data.

| Cluster | Genes |
| --- | --- |
| C1 | *Actb, Ctss, CT010467.1, Kctd12, Zfp36, Hexb, Rgs2, Sat1, Ivns1abp, Ddx5* |
| C2 | *Slc2a5, Adgrg1, Olfml3, Nav2, Ifngr1, Fscn1, Plcl1, Entpd1, Cmtm6, Tmem119* |
| C3 | *Csmd3, Jmjd1c, Nav2, Tanc2, Nav3, Ldlrad4, Dennd4a, Dock4, Csf1r, Zswim6* |
| C4 | *Cst3, Sparc, Csf1r, Selplg, Cx3cr1, Tmem119, Lgmn, Cmtm6, Slc2a5, Adgrg1* |
| C5 | *Itgb5, Selplg, Ier2, Csf1r, Eef2, Junb, Marcksl1, Psap, Fosb, Tmem119* |
| C6 | *Tmsb4x, Cst3, Rplp1, Eef1a1, Tpt1, Actb, Rps9, Rps3, Rps4x, Rpl32* |
| C7 | *Cdh23, Nkain2, Zfp36, Actb, Pde1c, Hspa1b, Kctd12, Wdr72, Mirg, Egr1* |
| C8 | *Fam189a1, Pde1c, Wdr72, Actb, Mirg, Sema3c, Ccl3, Ccl4, Serinc3, C1qa* |
| C9 | *Tanc2, Csmd3, Nav3, Nav2, Jmjd1c, Zfhx3, Srgap2, Rapgef5, Man1a, Btaf1* |
| C10 | *Cst7, Apoe, Gpnmb, Fth1, Spp1, Igf1, Lyz2, Clec7a, Ctsb, Lpl* |
| C11 | *Cst7, Apoe, Fth1, Spp1, Tyrobp, Lyz2, Ctsb, Gpnmb, Trem2, Eef1a1* |
| C12 | *Csf1r, Selplg, Ier2, mt-Cytb, Fosb, mt-Nd1, Fos, Itgb5, Lgmn, Marcks* |
| C13 | *Cst7, Gpnmb, Lpl, Apoe, Igf1, Itgax, Clec7a, Axl, Lgals3bp, Csf1* |
| C14 | *S100a6, Plac8, Ifitm6, Hp, Itgal, Cytip, Flna, Ifitm3, Lgals3, Cybb* |
| C15 | *Cst7, Spp1, Csf1, Ccl3, Apoe, Zfp36, H2-K1, Ccl4, Clec7a, Il1b* |
| C16 | *Il1r2, Saa3, Pla2g7, Chil3, S100a9, S100a8, Mt2, Lgals3, Msrb1, Apoe* |
| C17 | *Apoe, Itgax, Gpnmb, Cst7, Spp1, Csf1, Axl, Clic4, Lgals3bp, Osbpl8* |
| C18 | *Slc1a3, Clu, Aldoc, Plpp3, Sparcl1, Gpr37l1, Gja1, Mfge8, Cpe, Gpm6a* |
| C19 | *Nlgn1, Mgmt, Mirg, Fam189a1, Focad, Wdr72, Mcm4, Olfr1062, AC141639.4, Pde1c* |
| C20 | *Zc3h7a, Rgs1, Camk1d, Adam10, Ly6c1, Flt1, Epas1, Itm2a, Ly6a, Sptbn1* |
| C21 | *Catspere1, Nfasc, Myzap, Gm3383, Rab7, Slc15a2, Tsga10ip, Gm17501, Gm26917, Gm10406* |
| C22 | *Catspere1, Nkain2, Recql4, Slc5a1, Tsga10ip, Actb, Cdh23, Chl1, Dnm3, Camk2b* |
| C23 | *Gm12295, Fgf14, Zbtb16, Negr1, Lamc3, Cacna2d3, Myh11, 5330417C22Rik, Malrd1, Cplx2* |

